# Continuous Production of Recombinant Adeno-Associated Virus in the Insect Cell/Baculovirus Expression Vector System

**DOI:** 10.1101/2025.10.06.680831

**Authors:** John Joseph, Francesco Destro, Arella Yuan, Daniel Antov, Wenyu Chen, Sally Song, Matthew Theriault, Chengcheng Yuan, Alexander Sansom, Chiara Lundin, Tyler Burns, Caleb Neufeld, Jacqueline M. Wolfrum, Prasanna Srinivasan, Paul W. Barone, Anthony J. Sinskey, Richard D. Braatz, Stacy L. Springs

## Abstract

Continuous production processes may offer significant advantages for biotherapeutic manufacturing, including increased productivity, consistent product quality, reduced facility footprint, and decreased process turnaround time. Despite these benefits, the in-situ formation of defective recombinant baculovirus expression vectors (BEVs) hinders the continuous manufacturing of recombinant adeno-associated viruses (rAAV) in the baculovirus expression vector system. This study investigates an approach of reducing defective viruses through the infusion of standard recombinant baculovirus (rBV) and the compartmentalization of early- and late-stage infected cells, resulting in stable rAAV production. In this study, rAAVs were continuously produced in a series of cascading reactors, comprising a feeder reactor, an infection reactor, and a production reactor. Residence times and transfer rates across the three reactors were optimized based on the production kinetics of rBV and rAAV derived from our mechanistic model. The majority of rBV was produced within the production reactor, thereby reducing the impact of defective viruses in the infection reactor, enabling continuous rAAV production. This study demonstrates the successful implementation of a continuous rAAV production process, yielding over 5×10^10^ vg/mL per day for 4 weeks. This work represents the first reported continuous rAAV production process utilizing the Sf9/BEVS platform and establishes engineering know-how for overcoming manufacturing challenges associated with rAAV-based gene therapies.

## 1.0 Introduction

Recombinant adeno-associated virus (rAAV) has become the predominant vector for *in vivo* gene therapy, due to its non-pathogenic nature, broad tissue tropism, and prolonged gene expression^1–3^. These promising attributes paved the path to over 300 ongoing clinical trials using rAAVs for various conditions, ranging from metabolic diseases to cancers^4^. Despite clinical success, vector production with a consistent quality at a scale suitable for widespread use remains a major challenge^5^. As of July 2025, five approved gene therapies utilize rAAVs packaged in HEK293 (mammalian) cells, while three rely on the insect cell/baculovirus expression system (IC/BEVS)^6^. Given the extremely high dose requirements, ranging from 1.5×10^11^ to 2×10^11^ viral genomes (vg) for localized administration, and up to 1.33×10^14^ vg/kg for systemic delivery^7^, there is a growing demand for a scalable and cost-effective vector production process^8^.

The state-of-the-art of rAAV production is batch mode, either using IC/BEVS or triple-transient transfection of HEK293 cells. Although batch production is well-established, continuous manufacturing represents a promising avenue for cost reduction and enhanced quality assurance^9^. For instance, certain small molecules and biologics can be manufactured using continuous production methods, which may offer several advantages: (i) high product titer, (ii) reduced human intervention and risk of contamination, (iii) low variability between harvests, and (iv) lower cost^10,11^. Biologics manufactured using a continuous production mode include monoclonal antibodies, enzymes, and virus-like particles (VLPs)^12–14^. For monoclonal antibodies, it was estimated that continuous production could decrease 68% of manufacturing costs in clinical trials and 35% for commercial use^15^. In addition, monoclonal antibodies produced using a Chinese hamster ovary cell line in a continuous run over 5 weeks were both consistent in quality and therapeutic efficacy^12^. Similarly, enzymes such as PNPGase have been produced consistently with continuous runs of up to 40 days^13^. More recently, influenza VLPs were continuously generated through a production process using insect cells that lasted 20 days^14^. These previous endeavors indicate that continuous manufacturing could be a promising approach for IC/BEVS for rAAV production.

Production of rAAV in continuous mode is difficult to achieve using transient transfection in HEK293, producer cell lines, and IC/BEVS, due to several factors that adversely affect yield, as discussed below. Although transient transfection is employed in large-scale batch production, its implementation in continuous mode is hampered by the large dose of vector plasmids required for repeated transfection and inefficient plasmid uptake on repeated transfection^16,17^. In addition, the preparation of fresh polyplex complexes and repeated dosing add to the cost and introduce the risk of contamination. Another bottleneck to continuous operation is the cytostatic and cytotoxic nature of nonstructural Rep proteins, leading to negative selection pressure, which may result in deletion of genes from stable producer cell lines^18,19^. The IC/BEVS is an attractive alternative to HEK293 cells due to the higher yield of full rAAV capsids that can be achieved^20^. Further, the self-replication of recombinant baculovirus (rBV) in insect cells offers a natural avenue for cost reduction as infected cells continuously produce progeny rBV, eliminating the need for frequent manual intervention during vector production and reducing reliance on expensive GMP grade plasmid vectors required in HEK293-based systems^21^. However, a major issue associated with the rBV-based system is genetic instability, which complicates the maintenance of a consistent multiplicity of infection (MOI) of baculovirus with recombinant AAV genes^22^. Upon serial passaging, rBVs are prone to mutations that lead to the accumulation of two classes of defective viruses: (i) baculoviruses that have lost essential recombinant AAV genes, such as *rep, cap*, or the inverted-terminal-repeat-flanked gene of interest (GOI), and (ii), Defective Interfering Particles (DIPs) that lack critical genes required for baculovirus self-replication^23^. Both types of defective viruses disrupt the production of functional rAAV by impairing Rep and Cap protein expression, ultimately leading to a progressive decline and eventual halt of vector production.

In this study, we develop and quantify the first process for IC/BEVS-based continuous rAAV manufacturing that sustains high titers for four weeks. The process integrates a three-reactor cascade (growth, infection, production) with controlled infusion of low-passage recombinant baculovirus to separate cell proliferation, primary infection, and late-stage baculovirus budding and rAAV production. Process design is guided by serial-passage characterization of baculovirus stability and a mechanistic model of baculovirus infection and propagation that predicts the process outcome for diverse operating conditions through *in silico* simulation.

## 2. Results

### 2.1 Continuous rAAV production in the IC/BEVS fails due to rBV instability

A TwoBac system was used to evaluate continuous production of rAAV2/5 in insect cells (Figure 1a). The first type of rBV encodes the structural protein (cap5) and the AAV replication protein Rep2, while the second rBV carries the GOI flanked by ITR elements. A reporter gene, enhanced green fluorescent protein (eGFP), was used as the GOI to monitor transduction efficiency.

**Figure 1.**
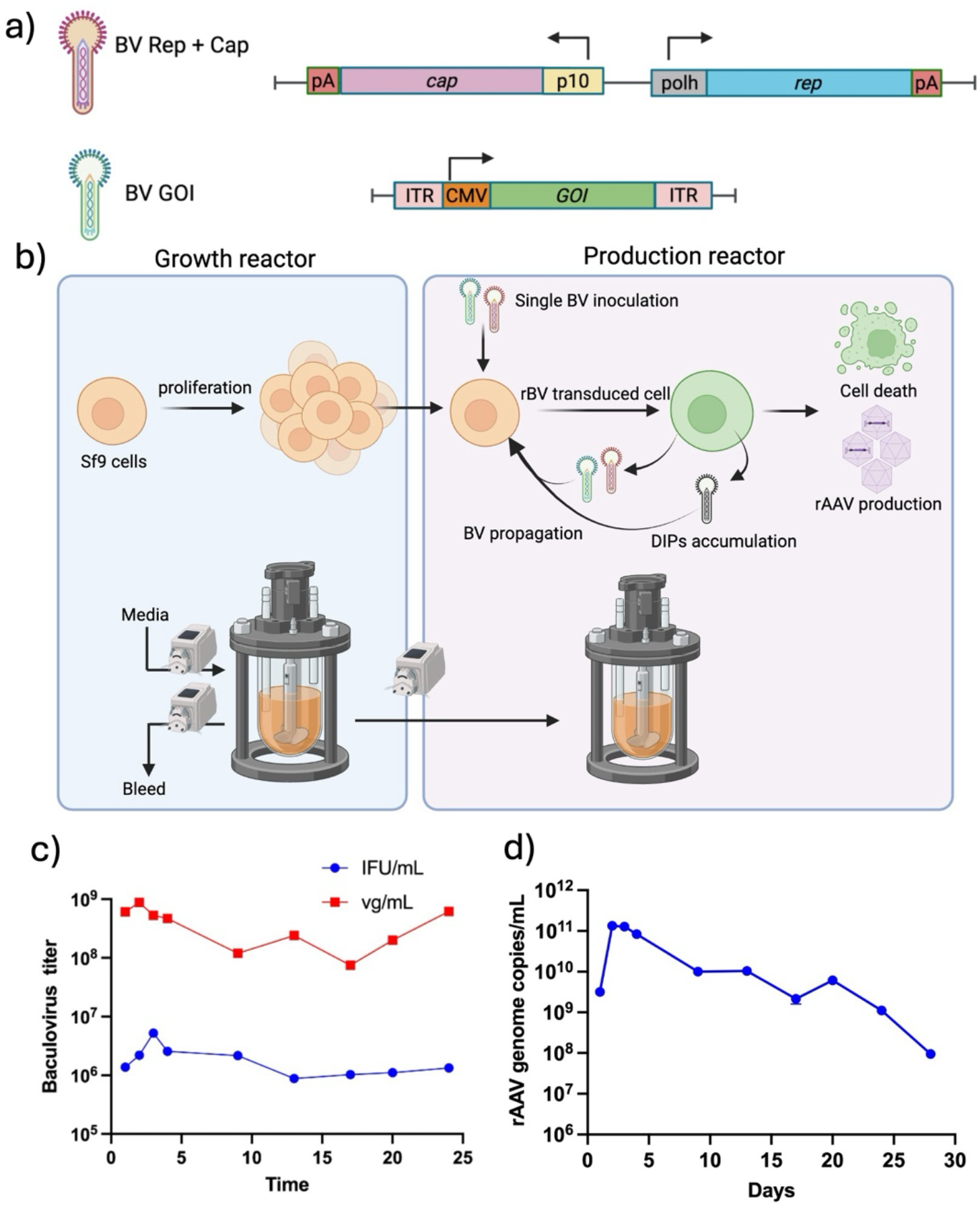
Continuous rAAV production in the IC/BEVS with a two-tank cascade. a) TwoBac system encoding GOI and AAV *rep* and *cap* genes for packaging rAAV in Sf9 cells. b) Two-reactor configuration with a residence time of 60 hours per tank, facilitating the growth of Sf9 cells in the growth reactor and the self-propagation of rBV in the production reactor. c) Budded baculovirus virions quantified from cell supernatant. Total baculovirus genome titer and infectious forming units per mL. d) Full rAAV titer quantified from cell suspension.

A two-reactor cascade, consisting of a growth reactor and a production reactor (Figure 1b), was implemented to enable continuous rAAV production over 4 weeks. Sf9 cells were maintained at a density of 2.5–4 million cells/mL in the growth reactor throughout the process. A single inoculation of the TwoBac system at an MOI of 2 plaque-forming unit (PFU) per cell in the production reactor initiated rBV self-replication and rAAV production. After 48 hours, uninfected Sf9 cells from the growth reactor were continuously transferred to the production reactor at a flow rate of 104 μL/min using a peristaltic pump. An average residence time of 60 hours was maintained in both reactors to support cell replication in the growth reactor and rAAV production in the production reactor. Samples collected from the production reactor and a downstream harvest tank during the continuous run were analyzed to quantify total BV genome and infectious titer along with rAAV genome, and capsid titer (Figures 1cd and S1).

The initial rAAV genome titer exceeded 1×10^11^ vg/mL post 72 hours rBV inoculation, but a sharp decline below 1 vg/mL occurred after 12–15 days. By days 22–25, the titer dropped below 1×10^9^ vg/mL, indicating that rAAV production had effectively reduced. Despite this decline, total and infective BV titers remained relatively stable throughout the process (Figure 1c). These results suggest the accumulation of baculoviruses lacking the *rep* and/or *cap* cassettes are likely contributing to the observed loss of rAAV production.

### 2.2 Analysis of rBV genetic stability via serial passage

The decline in rAAV production observed in the two-tank cascade system prompted further investigation into the genetic stability of the rBV vectors used in the process. A serial passage of rBV encoding *rep* and *cap* gene cassettes was performed to investigate the dynamics of mutant rBV accumulation during vector production. rBVs were sub-cultured every 72 hours at an MOI = 0.1 PFU/cell (Figure 2a). Primers were designed to target late expression factor 2 (*lef-2*) and rAAV gene (*rep* and *cap*) cassettes to quantify the total BV and rBV genome titers, respectively, using ddPCR. Loss of rAAV genes and protein expression was observed with increasing passage number of rBV (Figure 2b-e). Deletion of the recombinant genes from rBV generates defective baculoviruses that interfere with rAAV production. Due to the shorter genome, the defective particles may potentially replicate and outcompete rBVs during continuous cultivation.

**Figure 2.**
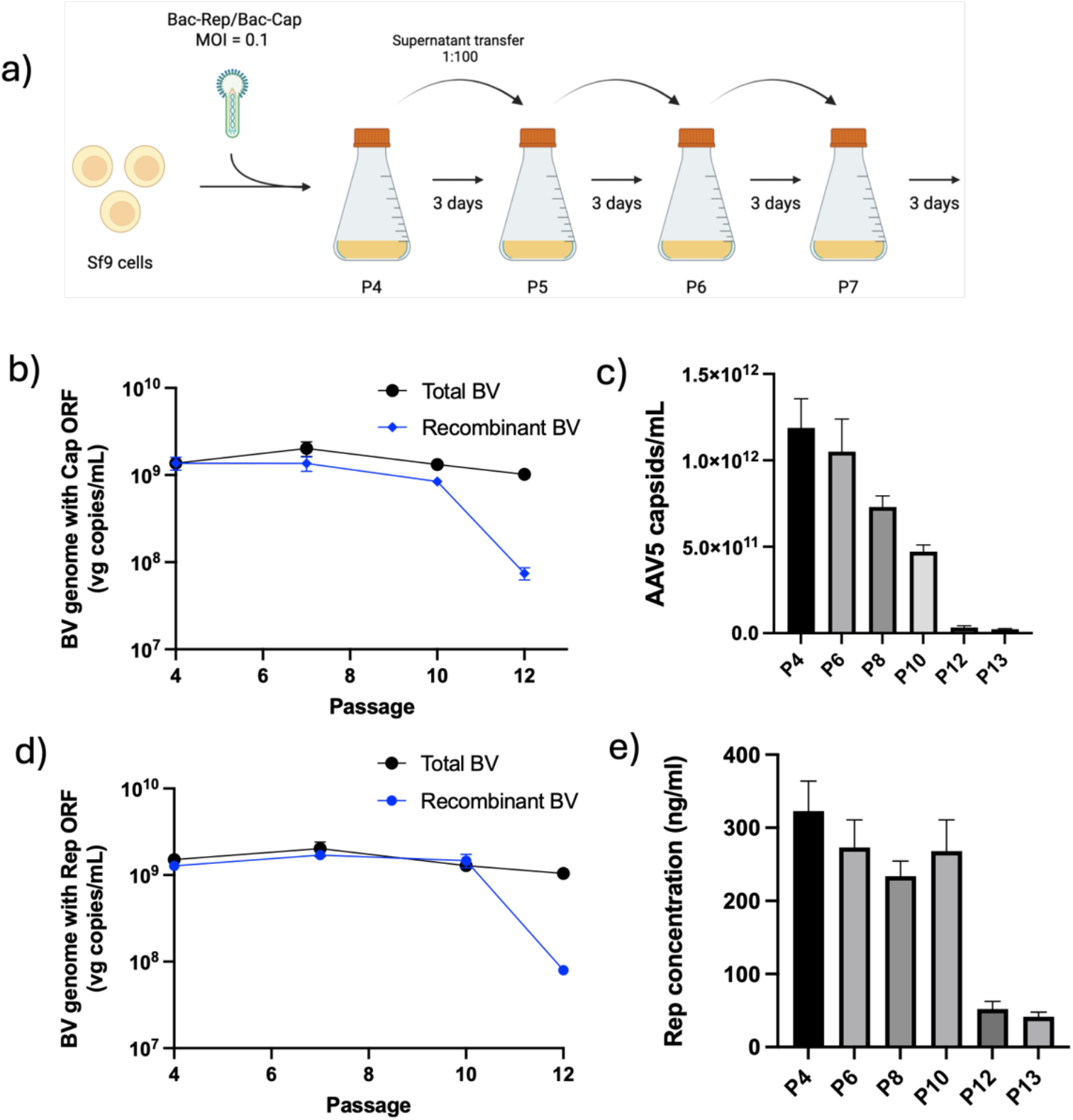
Analysis of genetic stability of rBV encoding rAAV genes through serial passages. a) Experimental layout of serial passage for rBV in Sf9 cells at MOI = 0.1. Cell culture supernatant containing the budded rBV is subcultured into a new flask containing uninfected Sf9 cells every 72 hours. b-e) Quantification of baculovirus and AAV proteins. b) & d) Total budded BV and rBV were quantified from cell supernatant using ddPCR. c) & e) Capsid and Rep proteins were quantified from cell suspension using ELISA.

### 2.3 Process design for continuous rAAV production in the IC/BEVS

The experiments of continuous rAAV production in the two-tank cascade and BV serial-passage showed that rAAV productivity declines when producer cells are predominantly infected by high-passage BVs lacking *rep* and/or *cap* cassettes. To mitigate this effect, a three-reactor cascade was designed, comprising growth, infection, and production bioreactors, with continuous cell transfer among reactors and continuous supplementation of low-passage (passage *N* = 4) rBV to the infection reactor (Figure 3a). This design aims at physically compartmentalizing (i) cell proliferation, (ii) primary infection, and (iii) baculovirus budding and rAAV production, leveraging the infection age dependency of baculovirus kinetics. Specifically, baculovirus reinfection becomes negligible beyond 3–5 hours post-infection (hpi) due to virus-induced receptor down-regulation^24^. In contrast, progeny baculovirus budding typically begins around 14–18 hpi and continues until cell lysis^24^. rAAV production is governed by very late promoters encoded in rBV that drive *rep* and *cap* expression, which are activated between 18–24 hpi^25^.

**Figure 3.**
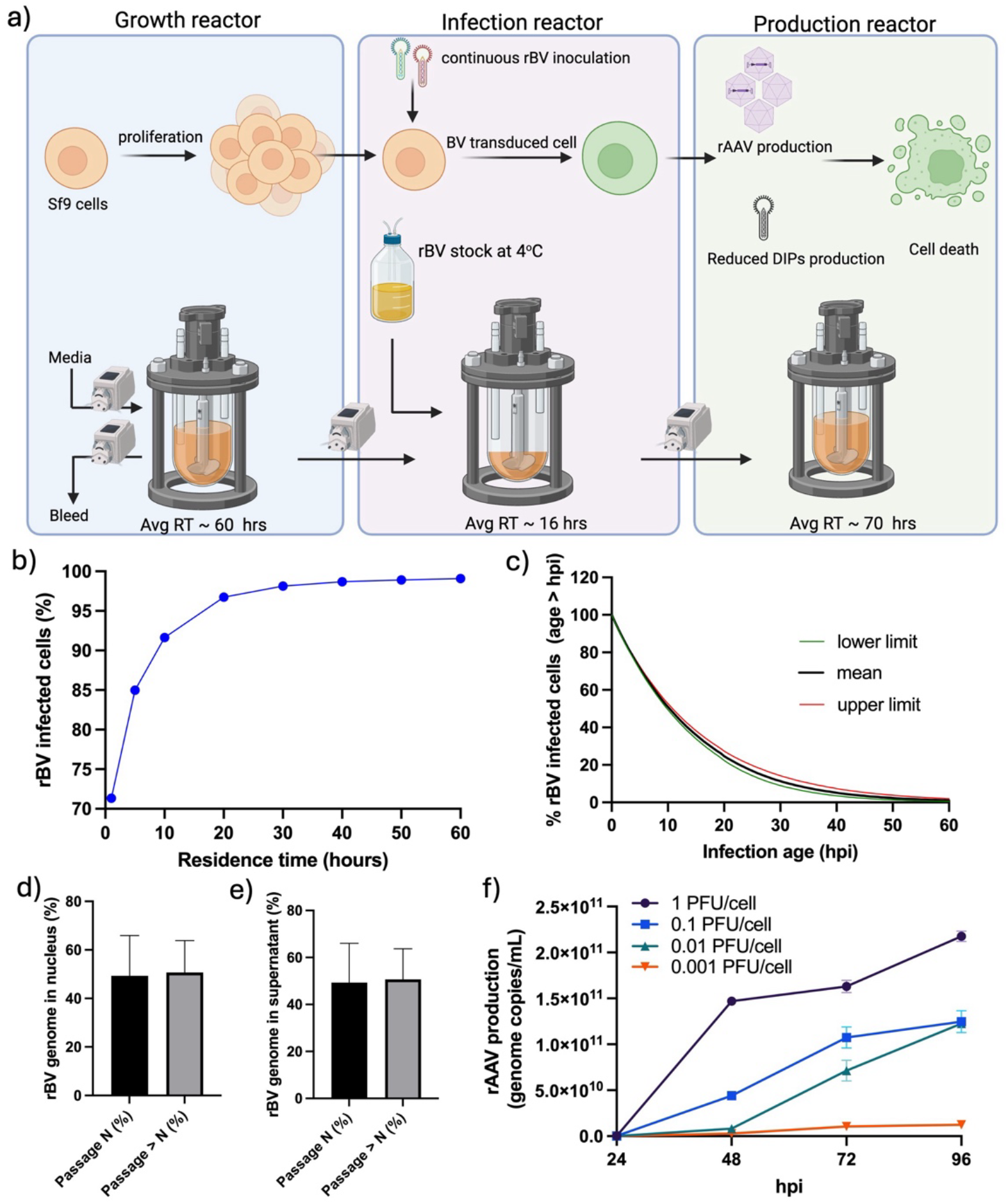
Process design for continuous rAAV production in the IC/BEVS with a three-tank cascade. a) Scheme of three-reactor configuration. b-e) Mechanistic model predictions for the infection reactor at steady-state, with a feed of 2.5 million uninfected cells/mL and 2 PFU/cell: b) Percentage of Sf9 cells infected with rBV as a function of residence time (RT), c) Infection age distribution of rBV-infected Sf9 cells for residence time equal to 16 hours. The y-axis indicates the percentage of infected cells with infection age greater than the indicated hpi based on lower limit, mean, and upper limit estimates d) & e) passage *N* vs. passage greater than *N* rBV copy number abundance in, respectively, the supernatant and nucleus of infected cells, for residence time equal to 16 hours. The model predictions shown in c–e are the mean predictions across 200 Monte Carlo realizations, with 95% confidence limits reflecting uncertainty propagated from the joint parameter distribution. f) rAAV production with the TwoBac platform at varying MOIs (0.001 to 1 PFU/cell per type of rBV) in a shake flask culture. The graph depicts the full rAAV genome copies determined using ddPCR from the cell lysate.

To implement this strategy effectively, a model-based process design was conducted to select appropriate values for the most critical operating conditions, namely the residence times of each reactor and the rBV concentration in the infection reactor feed. The objective was to ensure high infection rates from low-passage rBV in the infection reactor while minimizing reinfection in the production reactor, where defective baculoviruses accumulate. A mechanistic model of baculovirus infection and propagation in suspension cultures (Materials and Methods) supported the process design. The model enabled prediction of the infection-age distribution and the proportions of passage *N* and higher-passage baculoviruses in the infection reactor, both in the supernatant and within infected cells, under different operating conditions. Based on these simulations, a residence time of 16 hours, a feed of 2.5 million uninfected cells/mL, and 2 PFU/cell per rBV type were selected to ensure more than 95% of the cells in the infection reactor are infected at steady state (Figures 3b and S2). Figure 3c depicts the model prediction of the steady-state infection age distribution of infected cells in these operating conditions. More than 80% of infected cells exceed 3 hpi of infection age, thereby minimizing reinfection in the downstream production reactor (Figure 3c). Only 30% of infected cells exceed 18 hpi (onset of BV progeny budding), while 20% and 3% exceed 24 and 48 hpi, respectively. Under these conditions, the model simulations also indicate that passage *N* rBV constitutes more than 50% of the BV genomes in the supernatant, with infected cells similarly containing more than 50% of their total nuclear BV genome derived from passage *N* (Figures 3de).

Additional simulations were conducted to evaluate alternative process designs that were ultimately deemed less favorable. An infection reactor with the same feed (2.5 million uninfected cells/mL and 2 PFU/cell per rBV type) but a higher residence time resulted in the predominance of high-passage BVs at steady state (Figure S2). For example, a 60-hour residence time reduced the passage *N* genome fraction in infected cells and the supernatant to approximately 25% (Figure S3). This outcome underscores the advantage of a three-reactor system with a dedicated infection stage, which helps to maintain a high proportion of low-passage rBV in contrast to a simpler two-reactor configuration. Further simulations showed that supplying 1 PFU/cell of each rBV type in the feed maintains approximately 45% passage *N* genomes in infected cells (Figure S4). These findings support the selection of 1–2 PFU/cell as an effective feed condition for preserving a low-passage rBV population during continuous operation.

While maintaining low-passage rBV is essential to minimize the accumulation of defective particles, rAAV production also requires that individual cells are coinfected by both baculovirus types (encoding *rep, cap*, or the ITR-flanked GOI). To evaluate the relationship between MOI and productive coinfection, shake flask experiments were conducted to evaluate rAAV production across a range of MOIs for rBV encoding *rep* and *cap* genes to determine the MOI required in the infection reactor to achieve coinfection from both types of rBV and high rAAV production (Figure 3f). The results indicated that rAAV production increases with MOI, reaching a plateau at 2 PFU/cell, indicating that this level is sufficient to ensure coinfection in most cells. These experimental findings were further supported by a previously developed mechanistic model for rAAV production in the IC/BEVS, which indicated that MOIs of 2 PFU/cell led to productive coinfection and high rAAV yields in approximately 75% and 90% of cells, respectively, in shake flask cultures^26^. In the continuous process, the effective MOI in the infection reactor reflects both the baculovirus feed and a (limited) amount of progeny rBV generated *in situ*, which partially retains functional *rep* and *cap* genes.

Based on the experimental and modeling results, a residence time of 16 hours was selected for the infection reactor, with a target of 1.5–2 PFU/cell to simultaneously support productive coinfection and preserve a sufficient proportion of low-passage rBV. Residence times of 60 and 70 hours were chosen for the growth and production reactors, respectively, to allow sufficient time for cell proliferation and rAAV production, consistent with Sf9 growth and rAAV expression kinetics in the IC/BEVS.

### 2.4 Continuous, stable rAAV production in a three-tank cascade with continuous rBV feed

Continuous rAAV production in a three-tank cascade was implemented based on the process design described in the previous section (Figure 3a). Sf9 cells were maintained at a cell density of 2.5–4 million cells/mL in the three reactors during production. A constant feed of rBV was supplied via a peristaltic pump into the infection reactor, ensuring approximately 1.5–2 PFU/cell per type of rBV in the overall feed to the infection reactor. Samples from the infection reactor were collected to identify defective baculoviruses using ddPCR. To this end, we designed primers to target the *rep* and *cap* genes and *lef-2*. Based on the ddPCR readout, it is evident that all three primers showed similar gene amplification, which is indicative of *rep* and *cap* sequence conservation in the continuous production process (Figure 4a). The rAAV genome and capsid titers were quantified from the infection reactor, production reactor, and harvest during the continuous run for 28 days using ddPCR and ELISA (Figure 4bc). The rAAV genome and capsid titers showed stable production over the 28-day production period (Figure 4bc). Total rep proteins quantified from cells lysate shows stable expression during the production period (Figure 4d). SYPRO® Ruby staining of denatured rAAV capsids produced from the continuous production were performed to analyze VP1, VP2 and VP3 proteins (Figure 4e)

**Figure 4.**
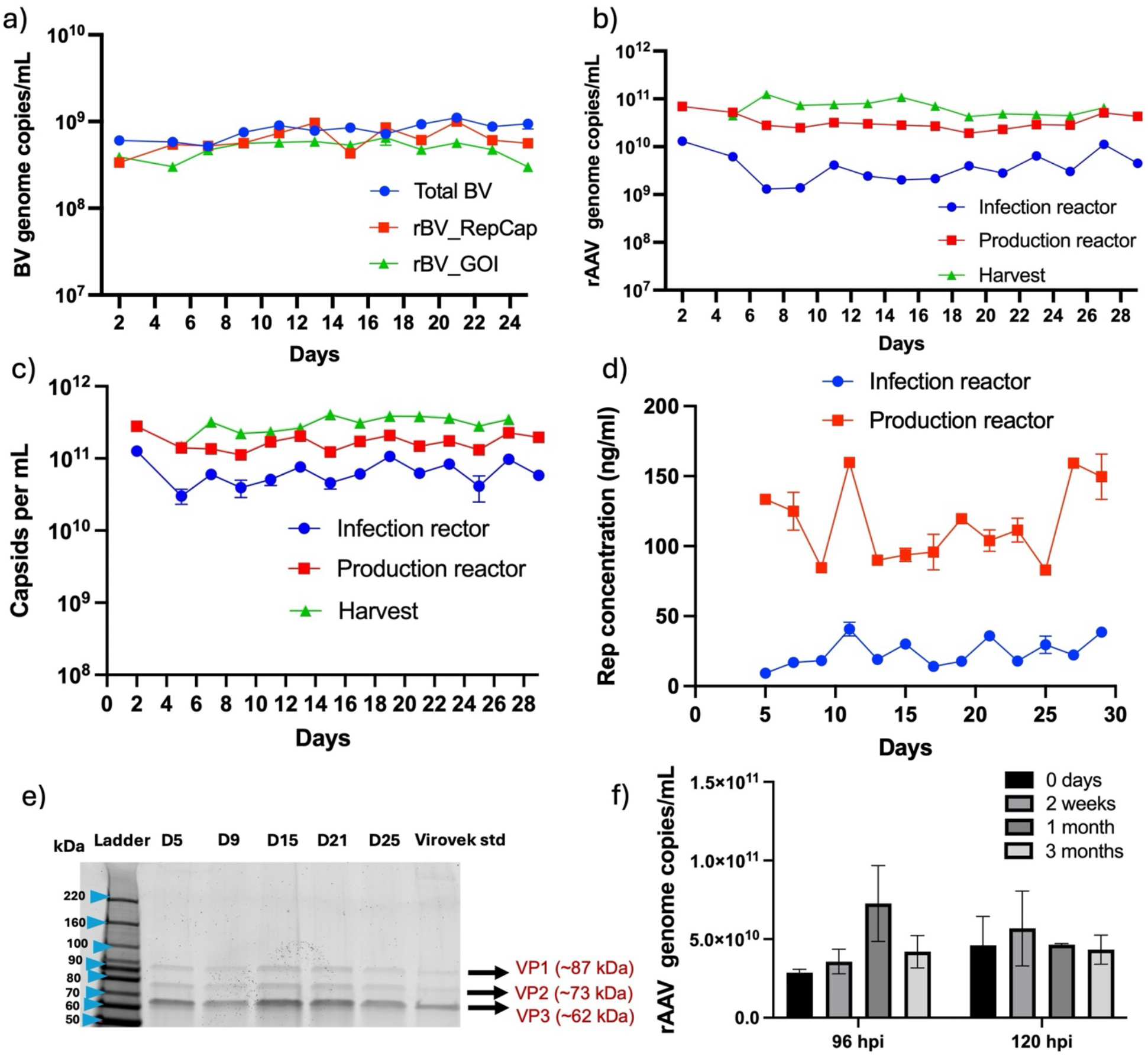
Continuous rAAV production using a three-tank cascade. a) Total BVs and rBVs were quantified from cell culture supernatant using ddPCR with primers to target the *lef2* and AAV genes b) rAAV genome titer was quantified from cell suspension quantified using ddPCR. c) quantification of total AAV capsid produced during the continuous production from cell suspension d) Rep concentration from cell lysates using ELISA e) SYPRO® Ruby staining of denatured AAV capsids produced from the continuous production showing viral proteins (VPs). f) Infectivity of budded rBV stored in spent media for 3 months at 4°C. The budded rBVs collected at different time points were used to infect Sf9 cells to produce rAAV.

The shelf stability of rBVs in crude spent media stored at 4°C was used for the rAAV production and showed no statistically significant difference (Figure 4f). Overall, the experiment yielded 4.5×10^14^ full rAAV capsids across 28 days, representing approximately 300% increase compared to the two-tank system control described in Section 2.1. Compared to batch-mode production over the same 28-day period, the process achieved an 80% increase in total rAAV output. This comparison assumes the same three bioreactors are used in batch mode for four weeks, with one batch per run per week, and a final full rAAV titer of 1×10^11^ vg/mL at the end of each batch.

An additional control experiment was carried out in the three-tank system with a 60-hour residence time for each reactor and no continuous rBV feed to the infection reactor (Figure S5). Sharp declines in the rAAV genome and capsid titer were registered 10 days after the process onset (Figure S5a, b). The rAAV genome titer decrease correlates with the reduced Rep protein expression (Figure S5c). In the next section, a fast and efficient approach that was employed to generate the baculovirus stock is described.

### 2.5 Fast, efficient generation of the baculovirus stock for continuous rBV feed

An efficient expansion step is required for producing a sizable viral stock to sustain continuous infusion of both types of rBV for 4 weeks. We developed an approach to expand the rBV stock and maintain rBVs at a low passage number for several weeks. In our study, 20 μL of rBV stock preserved as baculovirus-infected insect cells (BIICs) was scaled up to 400 mL in 5 days. rBVs were harvested using inline filters (1 micron pore size) post 5 days of infection with rBV at MOI= 0.0001 PFU/cell. ddPCR was performed to quantify rBV, and the ratio of both types of rBV was adjusted to 1:1 before storage at 4°C (Figure 5a). The production kinetics of rBV from the bioreactors revealed a rBV yield above 1×10^10^ vg/mL (Figure 5b). The rAAV production from budded rBV at different titers was compared to the rAAV production method using BIICs, as demonstrated in Figure S6. A viral titer above 10 BV/cell showed rAAV production comparable to BIICs at a dilution (1:100, approximately 1 PFU/cell).

**Figure 5.**
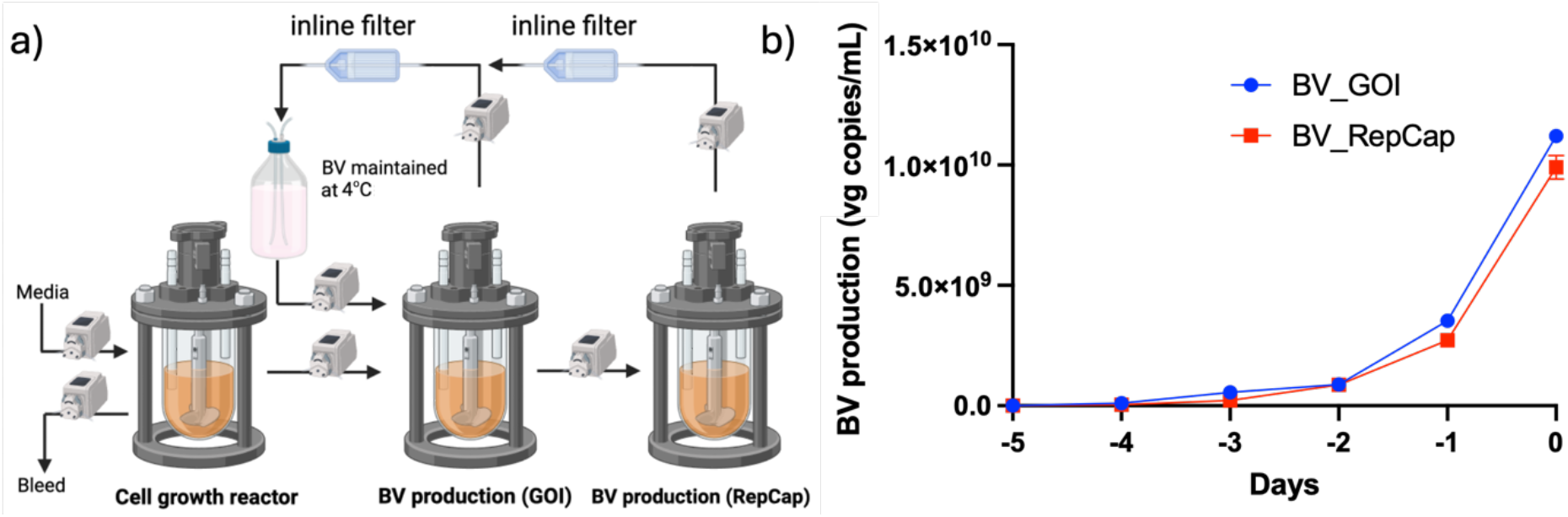
Protocol for generation of the recombinant baculovirus stock for continuous rAAV production. This involves the expansion of rBV from a single inoculation, followed by the harvest of budded rBV using inline filters. The two types of rBVs were quantified using ddPCR and combined to make the stock at a 1:1 ratio before storage at 4°C. b) Production kinetics and rBV volumetric yield of the budded virions during the stage 1 expansion step.

## 3. Discussion

Continuous rAAV production in the IC/BEVS is a promising strategy for reducing costs of goods sold and increasing volumetric productivity compared to traditional batch production. However, a significant challenge in implementing continuous production in the IC/BEVS is the formation and propagation of defective baculoviruses due to deletion events during rBV propagation^27^. Our first attempt of continuous rAAV production in the IC/BEVS, employing a two-reactor cascade, was unsuccessful, with rAAV yields declining significantly after 1–2 weeks of continuous operation (Figure 1).

Our serial passage experiments provided clear evidence of an increase in defective baculovirus populations lacking *rep* and/or *cap* genes, which became predominant beyond passage 12 (Figure 2). These observations highlighted the necessity of maintaining low passage rBV populations to ensure genetic stability and sustained rAAV production. While previous studies demonstrated improved genetic stability using FlashBac compared to Bac-to-Bac systems to generate the rBV stock^14,28,29^, the long term genetic stability of FlashBac is yet to be investigated..

Here, we introduced a novel process for rapid, efficient transition of batch-optimized rBVs to continuous production without the need for baculovirus redesign. Our process comprises a continuous three-tank cascade of growth, infection, and production reactors, with continuous infusion of low-passage rBV stocks into the infection reactor (Figure 3a). The physical separation of infection stages provides control over infection dynamics, improving Sf9 cell replication, baculovirus propagation, and ultimately, rAAV production. Crucially, defective baculoviruses accumulate primarily in the production bioreactor, where cells arrive at late infection stages and are refractory to reinfection. A mechanistic model of baculovirus infection and propagation supported the process design, enabling in silico simulations that evaluated infection efficiency and infection age and baculovirus passage distributions in the infection reactor, under varying residence times and rBV feed concentrations (Figures 3b–e and S2–4). Our experimental implementation validated these predictions, successfully demonstrating continuous rAAV production over a prolonged 28-day period.

We also developed a novel strategy for generating and maintaining low-passage rBV stocks throughout the whole duration of the continuous process. Notably, shelf stability studies demonstrated that rBV stocks retained their infectivity and showed no significant degradation after storage at 4°C for three months. This robustness enables practical storage flexibility, allowing the generation of the entire rBV stock several days before initiating continuous production without compromising productivity.

This study represents the first reported continuous rAAV production utilizing the IC/BEVS, effectively addressing critical manufacturing challenges related to defective baculovirus formation and genetic stability. Future process optimizations involving variables such as flow rate, cell concentration, and media formulation have the potential to further enhance yields. Moreover, the strategies and insights developed here can be broadly extended to other continuous viral manufacturing systems similarly challenged by defective virus formation.

### 4.0 Methods

Cell culture experiments, viral transduction, and quantification of viral titers were performed following BL2 procedures approved by MIT’s Environmental, Health and Safety (EHS) committee.

### 4.1 Cell lines and culture media

*Spodoptera frugiperda* (Sf9) suspension cells were maintained at a cell density ranging from 0.3 to 3×10^6^ cells/mL in serum-free SFM4 media (Hyclone Laboratories, Utah, USA). The Sf9 cells were cultured in a sterile 125-mL Erlenmeyer flask (Fisher brand sterile PC flasks, Cat. #PBV125). These cells were cultured at 27°C in an incubator (Thermo Scientific HERA cell VIOS 160i) on an orbital shaker (ORBI SHAKER™ CO2, Benchmark) at 135 rpm.

### 4.2 Serial passage of rBV

Sf9 cells, at a cell density of 1.5×10^6^ cells/mL, were added to a 125 mL Erlenmeyer flask and inoculated with recombinant BV encoding *rep* and *cap* genes at an MOI = 0.1 PFU/cell. Supernatant was harvested every 3 days and subsequently inoculated into a new flask of uninfected Sf9 cells at an MOI = 1 PFU/cell from passages 4 to 12. Samples collected from passages 4, 7, 10, and 12 were used for ddPCR after digestion with a DNAse I (New England Biolabs, MA, USA).

### 4.3 Quantification of vector genome

Droplet Digital Polymerase Chain Reaction (ddPCR) reaction mix was prepared with EvaGreen Supermix following the manufacturer’s protocol. Samples were diluted in DNase/RNase-Free Deionized Water (ThermoFisher Scientific, MA, USA). Droplet generation was performed using Droplet Generation Oil for EvaGreen on the QX200 Automated Droplet Generator. Thermal cycling was performed with the C100 Touch Thermal Cycler, using a heated lid at 105°C and a sample volume of 40 µL and a protocol as follows: enzyme activation at 95°C for 5 min, 40 cycles of denaturation at 95°C for 30 sec and annealing and extension at 60°C for 1 min, signal stabilization at 4°C for 5 min, and signal stabilization at 90°C for 5 min. Droplet reading was done with the QX200™ Droplet Reader (BioRad, MA, USA), and all plate sealing was done with the PX1 PCR plate sealer.

#### 4.3.1 Quantification of recombinant and total baculovirus genome titer

Baculovirus titer budded from infected cells was quantified from the spent media using primers that target the AAV gene (r*ep* and c*ap*) cassettes and a conserved baculovirus gene (*lef-2*). The primer sequences are:

Rep Forward: 5’-GCAGACAATGCGAGAGAATG-3’

Rep Reverse: 5’-CACGGGAAAGCACTCTAAAC-3’

Cap Forward: 5’-CATCGGCACCTTGGTTATTG-3’

Cap Reverse: 5’-CTACCGGAAAGCGGATAGAC-3’

lef-2 Forward: 5’-GAAGAAGCTGCGTAGTATGCC-3’

lef-2 Reverse: 5’-GTAGTTCTTCGGAGTGTGTTGC-3’

#### 4.3.2 Quantification of AAV genome titer

Cells were lysed with 1x lysis buffer (20% Tris HCl, 1% MgCl2, 5% Tween 20, 74% Milli-Q Water) and then spun down for 5 minutes at 10,000 rpm to remove cell debris. Host genomic DNA, baculovirus genome, and unpackaged AAV genome were digested with 10U DNase I (New England Biolabs, Massachusetts, USA) for 1 hour and heat-inactivated at 75°C for 10 minutes. ddPCR was performed as above using eGFP primers with sequences

eGFP Forward: 5’-GCAAAGACCCCAACGAGAAG-3’

eGFP Reverse: 5’-TCACGAACTCCAGCAGGACC-3’

### 4.4 Quantification of total rAAV capsids

Cells were lysed with 1x lysis buffer for 1 hour, then were spun down for 5 minutes at 10,000 rpm to remove cell debris. AAV5 Xpress ELISA kits (PROGEN, Pennsylvania, USA), as per the manufacturer’s protocol. Optical density (OD) was measured via a BioTek Synergy H1 Plate Reader (Agilent Technologies, Vermont, USA). Unknown concentrations were interpolated via a 4-parameter logistic fit (4PL) for capsid titer ELISAs.

### 4.5 Determination of total Rep proteins

Cell lysates were used for the quantification of total Rep proteins using AAV Rep ELISA kit (Cell Biolabs, California, USA). Unknown concentrations were interpolated with a linear standard curve.

### 4.6 Continuous AAV production

Sf9 cells were inoculated in a stirred‐tank bioreactor (OmniBRx Biotechnologies, NC, USA and Applikon, Holland, The Netherlands) connected by peristaltic pumps that allowed for continuous production. All three reactors (vessel volume 500 mL) were inoculated at 0.5×10^6^ cells/mL in 300 mL (unless specified) SFM4 Insect media (HyClone Laboratories, Utah, USA). Each reactor was held at 27°C, a pH of 6.1–6.5, with 70% dissolved oxygen maintained using a microsparger. Reactors were also equipped with three‐blade marine impellers rotating at 130 rpm. The media stock was protected from light and maintained at 4°C. All infections were performed using a dual baculovirus system encoding the Rep/Cap proteins and the GOI.

#### 4.6.1 Single baculovirus inoculation

When the concentration of Sf9 cells in all reactors reached 3×10^6^ cells/mL, baculovirus-infected insect cells (BIICs) were introduced into the infection reactor (reactor 2) at an MOI of 1 PFU/cell. The bioreactor was subsequently operated in batch mode for 48 hours, after which continuous production was initiated by activating the peristaltic pumps at a flow rate of 85 µL/min. Cells in the growth reactor (reactor 1) were maintained at a concentration below 4×10^6^ cells/mL through controlled bleeding. Cells were replenished when the cell density was below 2×10^6^ cells/mL. Samples were collected every other day from all reactors to be used for determining AAV and capsid titers.

#### 4.6.2 Continuous baculovirus inoculation

Continuous AAV production by BV inoculation strategy was done through 2 stages:

##### 4.6.2.1 Baculovirus expansion in bioreactor

When SF9 cells in reactor 2 and reactor 3 reached 1.5×10^6^ cells/mL, both were inoculated with baculovirus at an MOI = 0.001 PFU/cell. GOI baculovirus was introduced into reactor 2, while Rep/Cap baculovirus was introduced into reactor 3. After infection, inline filters (Hyclone) were used to remove cellular debris from the cultures, and the clarified filtrate was pumped into a collection container maintained at 4°C.

##### 4.6.2.2 Continuous rBV feed

The clarified baculovirus-containing filtrate was continuously pumped into reactor 2 to maintain an MOI of 2 PFU/cell. After 24 hours, continuous AAV production was initiated by activating the peristaltic pumps, allowing for steady-state infection and capsid production.

### 4.7 Long-term stability of recombinant baculovirus

Sf9 cells were cultured at 1.5×10^6^ cells/mL with rBV infected cells at a dilution level of 1/10000 that had been preserved at –196°C. These cells were preserved in a freezing media consisting of 10% DMSO and 150 mM Trehalose in SFM4 Insect media (HyClone Laboratories, Utah, USA). Cells were checked daily until viability dropped below 80% and average cell diameter increased by 3–4 µL, indicating infection. The supernatant, collected after spinning down at 10,000 RPM for 5 minutes, was collected and stored at 4°C. Approximately every two days, samples were taken from this stored media containing the budded rBVs and added to media containing uninfected Sf9 cells (1.5×10^6^ cells/mL) at an MOI = 1 PFU/cell. The BV titer of the supernatant was measured using ddPCR as described above. After 96 hours, the cells were harvested and lysed. Cell lysis was performed as described above, and cell lysate was assayed for copies of both BV and AAV genomes via ddPCR, as well as capsid titer and Rep protein concentration via ELISAs.

### 4.8 Quantification of VP proteins ratio

Sf9 cell suspension was first lysed as detailed above, and cell lysate was purified using Dynabeads™ CaptureSelect™ AAVX Magnetic Beads (Thermo Scientific, Massachusetts, USA), as per the manufacturer’s protocol. Purified capsid samples were mixed with Laemmli SDS sample buffer, reducing (6x) (Thermo Scientific, Massachusetts, USA) and placed upon a heat block at 90°C. After 20 minutes, samples, along with BenchMark™ Protein Ladder (Invitrogen, Massachusetts, USA), were loaded onto a gel, and sodium dodecyl sulfate-polyacrylamide gel electrophoresis (SDS-PAGE) was performed. 4–15% Mini-PROTEAN^®^ TGX™ Precast Protein Gels (Bio-Rad, Massachusetts, USA) and 1x Tris/Glycine/SDS (Bio-Rad) were used for the electrophoresis. Gel was then stained with SYPRO^®^ Ruby Protein Gel Stain (Invitrogen, Massachusetts, USA) according to the manufacturer’s protocol. Images were taken using a ChemiDoc MP Imaging System (Bio-Rad).

### 4.9 Mechanistic model of baculovirus infection and propagation

The mechanistic model of baculovirus infection and propagation is adapted from Destro and Braatz, where a more detailed description of the model is provided^30^. The model describes viral infection and propagation in a well-mixed tank, operated in batch or continuous mode. The model accounts for the presence of two viral species in the system: baculovirus of passage *N* (virus 1) and baculovirus of passage greater than *N* (virus 2). Based on the experimental conditions followed in this work, *N* = 4 was set. The full set of model equations is reported in the Supporting Information. Briefly, the model inputs are the initial conditions for all the system states (Table S1), the reactor operating mode (batch or continuous) and, for continuous operating mode, the residence time and the feed concentrations of viable and nonviable uninfected cells and virions of type 1 and 2. The model outputs are the time evolution profiles of all the system states, namely the concentration within the reactor volume of uninfected viable cells, viable cells infected only by virus 1 or 2, viable cells coinfected by virus 1 and virus 2, nonviable cells, free virions of type 1 and 2, virus 1 bound to viable cells infected only by virus 1, virus 2 bound to viable cells infected only by virus 2, virus 1 and virus 2 bound to viable coinfected cells, virus 1 in nucleus of viable cells infected only by virus 1, virus 2 in nucleus of viable cells infected only by virus 2, virus 1 and virus 2 in nucleus of viable coinfected cells. All concentrations of infected cells, cell-bound virus, and nuclear viral genomes are dynamically tracked over time and resolved by the relevant infection age(s): for cells infected by only virus 1 or only virus 2, the model tracks the corresponding infection age; for coinfected cells, both infection ages are tracked independently. The main viral infection and propagation steps captured by the model are: baculovirus infection, trafficking to nucleus, replication, and viral progeny production. The model explicitly accounts for the dependence of re-infection, intracellular replication, infected cell death, and virion release kinetics on the infection age of each cell. To generate the results presented in this manuscript, model inputs were specified according to the conditions of each experiment. The model equations were solved using a numerical method described previously^30^. All parameter values used in the simulations are listed in Table S2. These parameters, along with their confidence intervals, were originally estimated in Destro et al.^31^ based on a combination of published data and in-house experiments on baculovirus infection and propagation kinetics. The only parameter re-estimated for this study was the baculovirus progeny production rate, which was inferred via maximum likelihood estimation using data from the serial passage experiments. Uncertainty was propagated into the model predictions using 200 Monte Carlo realizations, each sampling a parameter set independently from the joint parameter distribution.

## Acknowledgements

The research was supported by the U.S. Food and Drug Administration under Contract No. 75F40121C00131. The Sf9 cells and recombinant baculoviruses encoding AAV genes were generously gifted by Robert Kotin, University of Massachusetts, USA. The authors gratefully acknowledge OmniBRx Biotechnologies, specifically Ravindra Patel, Anandprakash Joshi, Ravikumar Daraji for offering the bioreactors and technical support.

## Author contributions

John Joseph: Conceptualization; methodology; formal analysis; investigation; data curation; writing (original draft); writing (review and editing); visualization. Francesco Destro: Conceptualization; methodology; software; model validation; formal analysis; investigation; data curation; writing (original draft); writing (review and editing). Arella Yuan, Daniel Antov, Wenyu Chen, Sally Song, Matthew Theriault, Chengcheng Yuan, Alexander Sansom, Chiara Lundin, Tyler Burns: Methodology; investigation; Data curation; formal analysis; investigation; data curation; writing (review and editing), Caleb Neufeld: funding acquisition; Jacqueline M. Wolfrum: Formal analysis; resources; data curation; writing (review and editing); funding acquisition. Prasanna Srinivasan: Conceptualization; methodology; formal analysis visualization; investigation; Paul W. Barone: Conceptualization; methodology; formal analysis; resources; data curation; writing (original draft); writing (review and editing); funding acquisition. Anthony J. Sinskey: Resources; data curation; writing (original draft); writing (review and editing); supervision; funding acquisition. Richard D. Braatz: Conceptualization; methodology; resources; data curation; writing (original draft); writing (review and editing); funding acquisition. Stacy L. Springs: Conceptualization; methodology; resources; data curation; resources; writing (original draft); writing (review and editing); funding acquisition. All authors have reviewed and approved the final version of the manuscript.

