## Supporting information for "Continuous Production of Recombinant Adeno-Associated Virus in the Insect Cell/Baculovirus Expression Vector System"

### Additional information on the mechanistic model of baculovirus infection and propagation

The system states are summarized in Table S1. The description of all model parameters is reported in Table S2. The model equations for the concentrations of cellular species are

$$\frac{dT}{dt} = \mu T - k_{b,T}T(V_1 + V_2) - k_{d,T}T + D(T_{\text{in}} - T)$$

$$\frac{\partial i_1}{\partial t} + \frac{\partial i_1}{\partial \tau_1} = k_{b,T}TV_1\delta(\tau_1) - i_1\left(\bar{k}_{d,I_1} + D + \bar{k}_{b,I_1}V_2\right)$$

$$\frac{\partial i_2}{\partial t} + \frac{\partial i_2}{\partial \tau_2} = \mathrm{k}_{b,T}TV_2\delta(\tau_2) - i_2\left(\bar{k}_{d,I_2} + D + \bar{k}_{b,I_2}V_1\right)$$

$$\frac{\partial c}{\partial t} + \frac{\partial c}{\partial \tau_1} + \frac{\partial c}{\partial \tau_2} = \bar{k}_{b,I_2}i_2V_1 + \bar{k}_{b,I_1}i_1V_2 - c\left(\bar{k}_{d,C} + D\right)$$

$$\frac{dW}{dt} = k_{d,T}T + \int_0^\infty \bar{k}_{d,I_1}\, i_1\, d\tau_1 + \int_0^\infty \bar{k}_{d,I_2}\, i_2\, d\tau_2 + \int_0^\infty\int_0^\infty \bar{k}_{d,c}\, c\, d\tau_1 d\tau_2 - WD,$$

where $D$ is the dilution rate (equal to the reciprocal of the residence time), $\tau_1$ is the infection age with respect to virus 1, $\tau_2$ is the infection age with respect to virus 2, $\delta(\cdot)$ is the Dirac delta function, the equivalent binding kinetic parameters are calculated based on the cell infection age as

$$\bar{k}_{b,I_j}(\tau_j) = \begin{cases} k_{b,T}, & \text{for } \tau_j < \tau_b \\ k_{b,T}\exp\left(-\beta_b(\tau_j - \tau_b)\right), & \text{for } \tau_j \geq \tau_b \end{cases}, \text{for } j = \{1,2\}$$

$$\bar{k}_{b,C}(\tau_1,\tau_2) = \begin{cases} k_{b,T}, & \text{for } \max\{\tau_1,\tau_2\} < \tau_b \\ k_{b,T}\exp\left(-\beta_b(\max\{\tau_1,\tau_2\} - \tau_b)\right), & \text{for } \max\{\tau_1,\tau_2\} \geq \tau_b \end{cases}$$

and the equivalent death kinetic parameters are calculated as

$$\bar{k}_{d,I_j}(\tau_j) = \begin{cases} k_{d,T}, & \text{for } \tau_j < \tau_b \\ k_{d,T}\ln\left(\frac{n_{I_j}}{i_j}\right), & \text{for } \tau_j \geq \tau_b \end{cases}, \text{for } j = \{1,2\}$$

$$\bar{k}_{d,C}(t,\tau_1,\tau_2) = \begin{cases} k_{d,T}, & \text{for } \max\{\tau_1,\tau_2\} < \tau_d \\ k_{d,T}\ln\left(\frac{n_{C,V_1}(t,\tau_1,\tau_2) + n_{C,V_2}(t,\tau_1,\tau_2)}{c(t,\tau_1,\tau_2)}\right), & \text{for } \max\{\tau_1,\tau_2\} \geq \tau_d \end{cases}.$$

The model equations for the free virion concentrations assume that passage $N$ baculovirus (virus 1) is present in the feed with concentration $V_{1,\text{in}}$ and that all infected cells produce only baculovirus of passage greater than $N$ (virus 2):

$$\frac{dV_1}{dt} = -\int_0^\infty V_1\bar{k}_{b,I_1}\, i_1\, d\tau_1 - \int_0^\infty V_1\bar{k}_{b,I_2} i_2\, d\tau_2 - \int_0^\infty \int_0^\infty V_1\bar{k}_{b,C} c\, d\tau_1 d\tau_2$$
$$- V_1\left(k_{b,T}T + k_{d,V} + D - V_{1,\text{in}}D\right)$$

$$\frac{dV_2}{dt} = \int_0^\infty \left(\bar{k}_{v,I_1} - V_2\bar{k}_{b,I_1}\right) i_1\, d\tau_1 + \int_0^\infty \left(\bar{k}_{v,I_2} - V_2\bar{k}_{b,I_2}\right) i_2\, d\tau_2$$
$$+ \int_0^\infty \int_0^\infty \left(\bar{k}_{v,C} - V_2\bar{k}_{b,C}\right) c\, d\tau_1 d\tau_2 - V_2\left(k_{b,T}T + k_{d,V} + D\right).$$

The equivalent kinetic parameters for progeny release account for the infection-age dependency of baculovirus budding are

$$\bar{k}_{v,I_j}(\tau_j) = \begin{cases} 0, & \text{for } \tau_j < \tau_v^{\text{on}} \vee \tau_j > \tau_v^{\text{off}} \\ k_v, & \text{for } \tau_v^{\text{on}} \leq \tau_j \leq \tau_v^{\text{off}} \end{cases}, \text{for } j = \{1,2\}$$

$$\bar{k}_{v,C}(\tau_1,\tau_2) = \begin{cases} 0, & \text{for } \max\{\tau_1,\tau_2\} < \tau_v^{\text{on}} \vee \max\{\tau_1,\tau_2\} > \tau_v^{\text{off}} \\ k_v, & \text{for } \tau_v^{\text{on}} \leq \max\{\tau_1,\tau_2\} \leq \tau_v^{\text{off}} \end{cases}$$

The model equations for the concentrations of virus bound to cells and of viral genomes in the nucleus of infected cells are

$$\frac{\partial b_{I_1}}{\partial t} + \frac{\partial b_{I_1}}{\partial \tau_1} = V_1\left(k_{b,T} T\, \delta(\tau_1) + \bar{k}_{b,I_1} i_1\right) - b_{I_1}\left(\bar{k}_{d,I_1} + rD + \bar{k}_{b,I_1} V_2 + k_i\right)$$

$$\frac{\partial b_{I_2}}{\partial t} + \frac{\partial b_{I_2}}{\partial \tau_2} = V_2\left(k_{b,T} T\, \delta(\tau_2) + \bar{k}_{b,I_2} i_2\right) - b_{I_2}\left(\bar{k}_{d,I_2} + rD + \bar{k}_{b,I_2} V_1 + k_i\right)$$

$$\frac{\partial b_{C,V_1}}{\partial t} + \frac{\partial b_{C,V_1}}{\partial \tau_1} + \frac{\partial b_{C,V_1}}{\partial \tau_2} = \bar{k}_{b,I_1} V_2 b_{I_1} \delta(\tau_2) + \bar{k}_{b,I_2} V_1 i_2\, \delta(\tau_1) + \bar{k}_{b,C} c V_1 - b_{C,V_1}\left(\bar{k}_{d,C} + rD + \bar{k}_{i,C}\right)$$

$$\frac{\partial b_{C,V_2}}{\partial t} + \frac{\partial b_{C,V_2}}{\partial \tau_1} + \frac{\partial b_{C,V_2}}{\partial \tau_2} = \bar{k}_{b,I_2} V_2 b_{I_2}\, \delta(\tau_1) + \bar{k}_{b,I_1} V_2 i_1\, \delta(\tau_2) + \bar{k}_{b,C} c V_2 - b_{C,V_2}\left(\bar{k}_{d,C} + rD + \bar{k}_{i,C}\right)$$

$$\frac{\partial n_{I_1}}{\partial t} + \frac{\partial n_{I_1}}{\partial \tau_1} = \eta k_i b_{I_1} + \bar{k}_{r,I_1} n_{I_1} - n_{I_1}\left(\bar{k}_{d,I_1} + k_{d,N} + rD + \bar{k}_{b,I_1} V_2\right)$$

$$\frac{\partial n_{I_2}}{\partial t} + \frac{\partial n_{I_2}}{\partial \tau_2} = \eta k_i b_{I_2} + \bar{k}_{r,I_2} n_{I_2} - n_{I_2}\left(\bar{k}_{d,I_2} + k_{d,N} + rD + \bar{k}_{b,I_2} V_1\right)$$

$$\frac{\partial n_{C,V_1}}{\partial t} + \frac{\partial n_{C,V_1}}{\partial \tau_1} + \frac{\partial n_{C,V_1}}{\partial \tau_2} = \eta k_i b_{C,V_1} + \bar{k}_{r,C}\, n_{C,V_1} + \bar{k}_{b,I_1} n_{I_1} V_2 \delta(\tau_2) - n_{C,V_1}\left(\bar{k}_{d,C} + k_{d,N} + rD\right)$$

$$\frac{\partial n_{C,V_2}}{\partial t} + \frac{\partial n_{C,V_2}}{\partial \tau_1} + \frac{\partial n_{C,V_2}}{\partial \tau_2} = \eta k_i b_{C,V_2} + \bar{k}_{r,C}\, n_{C,V_2} + \bar{k}_{b,I_2} n_{I_2} V_1 \delta(\tau_1) - n_{C,V_2}\left(\bar{k}_{d,C} + k_{d,N} + rD\right),$$

Where the equivalent kinetic parameters for viral replication are

$$\bar{k}_{r,I_j}(\tau_j) = \begin{cases} 0, & \text{for } \tau_j < \tau_r^{\text{on}} \vee \tau_j > \tau_r^{\text{off}} \\ k_r, & \text{for } \tau_r^{\text{on}} \le \tau_j \le \tau_r^{\text{off}} \end{cases}, \text{for } j = \{1,2\}$$

$$\bar{k}_{r,C}(\tau_1, \tau_2) = \begin{cases} 0, & \text{for } \max\{\tau_1, \tau_2\} < \tau_r^{\text{on}} \vee \max\{\tau_1, \tau_2\} > \tau_r^{\text{off}} \\ k_r, & \text{for } \tau_r^{\text{on}} \le \max\{\tau_1, \tau_2\} \le \tau_r^{\text{off}} \end{cases}$$

**Table S1.** States of the mechanistic model of baculovirus infection and propagation. Virus 1 represents low-passage baculovirus (passage $N$, i.e., the passage supplied in the feed), whereas virus 2 denotes higher-passage baculovirus (passage > $N$) arising during propagation.

| Symbol | UOM | Description |
|---|---|---|
| $b_{I_j}(t, \tau_j)$ | virus mL$^{-1}$ hpi $^{-1}$ | Virus $j$ attached to $i_j(t, \tau_j)$, for $j = \{1,2\}$ |
| $b_{C,V_j}(t, \tau_1, \tau_2)$ | virus mL$^{-1}$ hpi $^{-2}$ | Virus $j$ attached to $c(t, \tau_1, \tau_2)$, for $j = \{1,2\}$ |
| $c(t, \tau_1, \tau_2)$ | cell mL$^{-1}$ hpi $^{-2}$ | Coinfected cells |
| $i_j(t, \tau_j)$ | cell mL$^{-1}$ hpi $^{-1}$ | Cells infected by virus $j$, for $j = \{1,2\}$ |
| $n_{I_j}(t, \tau_j)$ | vg mL$^{-1}$ hpi $^{-1}$ | Viral genome $j$ in nucleus of $i_j(t, \tau_j)$, for $j = \{1,2\}$ |
| $n_{C,V_1}(t, \tau_1, \tau_2)$ | vg mL$^{-1}$ hpi $^{-2}$ | Viral genome j in nucleus of $c(t, \tau_1, \tau_2)$, for $j = \{1,2\}$ |
| $T(t)$ | cell mL$^{-1}$ | Uninfected cells |
| $V_j(t)$ | virus mL$^{-1}$ | Extracellular virus $j$, for $j = \{1,2\}$ |
| $W(t)$ | cell mL$^{-1}$ | Nonviable cells |

**Table S2.** Parameters of the mechanistic model of baculovirus infection and propagation.

| Symbol | UOM | Value | 95% Confidence Interval | Description | Source |
|---|---|---|---|---|---|
| Infection | | | | | |
| $k_{b,T}$ | mL $cell^{-1}$ $h^{-1}$ | $6.3\times10^{-7}$ | ($5.2\times10^{-7}$, $7.4\times10^{-7}$) | Viral binding kinetic constant | Destro et al. (2023) |
| $\tau_b$ | hpi | 1.80 | (1.3, 2.3) | Onset of viral binding decay for infected cells | Destro et al (2023); Rohrmann (2019) |
| $\beta_b$ | [–] | 0.5 | – | Coefficient for viral binding decay for infected cells | Destro and Braatz (2024) |
| Cell growth and death | | | | | |
| $\mu$ | $h^{-1}$ | $2.8\times10^{-2}$ | ($2.7\times10^{-2}$, $2.9\times10^{-2}$) | Growth kinetic constant for uninfected cells | Mena et al. (2010); Power et al. (1994) |
| $k_{d,T}$ | $h^{-1}$ | $6.3\times10^{-7}$ | ($6\times10^{-5}$, $1\times10^{-4}$) | Death kinetic constant for uninfected cells | Power et al. (1994); Rohrmann (2019) |
| $k_{d,I}$ | $h^{-1}$ | $2.9\times10^{-3}$ | ($2.9\times10^{-3}$, $1\times10^{-4}$) | Death kinetic constant for infected cells | Destro et al. (2023) |
| $\tau_d$ | hpi | 24 | (22, 26) | Onset of death rate increase for infected cells | Destro et al. (2023); Rohrmann (2019) |
| Viral degradation | | | | | |
| $k_{d,V}$ | $h^{-1}$ | $7\times10^{-3}$ | ($1\times10^{-3}$, $1.3\times10^{-2}$) | Degradation kinetic constant for virions | Power et al. (2024) |
| $k_{d,N}$ | $h^{-1}$ | 0 | – | Degradation kinetic constant for nuclear viral genome | Destro et al (2023) |
| Viral trafficking and replication | | | | | |
| $k_i$ | $h^{-1}$ | 0.6 | (0.48, 0.72) | Virus internalization kinetic constant | Dee and Shuler (1997) |
| $\eta$ | [–] | 0.50 | (0.45, 0.55) | Fraction of internalized virus that reaches the nucleus | Dee and Shuler (1997) |
| $k_r$ | $h^{-1}$ | 0.732 | (0.651, 0.813) | Viral genome replication kinetic constant | Destro et al (2023) |
| $\tau_r^{\text{on}}$ | hpi | 6 | (5, 7) | Onset of viral genome replication in infected cells | Destro et al (2023); Rohrmann (2019) |
| $\tau_r^{\text{off}}$ | hpi | 18 | (17, 19) | End of viral genome replication in infected cells | Destro et al (2023); Rohrmann (2019) |
| Progeny release | | | | | |

| | | | | | |
|---|---|---|---|---|---|
| $k_v$ | PFU cell h$^{-1}$ | 0.65 | (0.5, 1.5) | Progeny release rate | Estimated from serial passage experiments data (Figure 3f) |
| $\tau_v^{\mathrm{on}}$ | hpi | 18 | (16, 20) | Onset of progeny release for infected cells | Destro et al (2023); Rohrmann (2019) |
| $\tau_v^{\mathrm{off}}$ | hpi | Cell death | – | End of progeny release for infected cells | Destro et al (2023); Rohrmann (2019) |

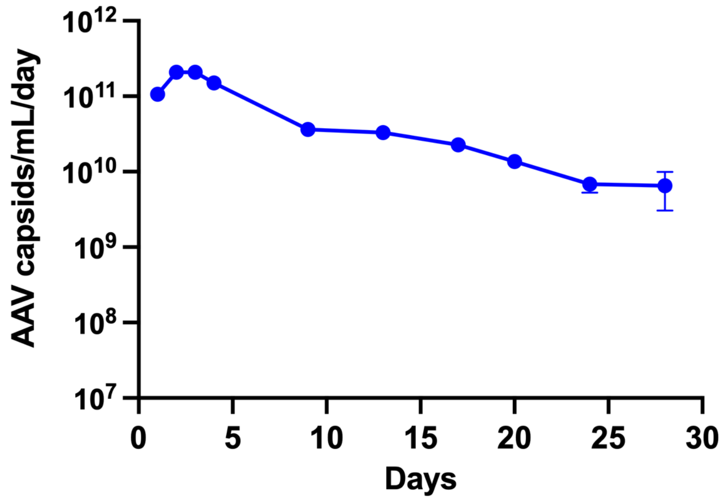


**Figure S1.** Total capsid titer in the experiment of continuous rAAV production in the IC/BEVS with a two-tank cascade (residence time of 60 hours per tank and no continuous rBV feed).

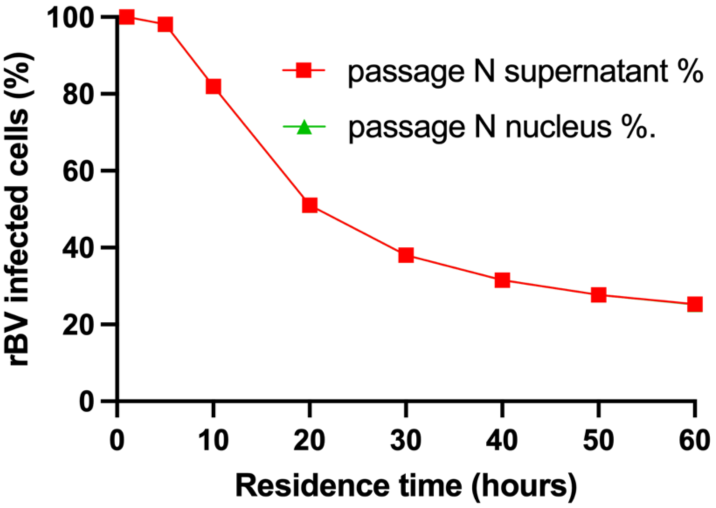


**Figure S2.** Mechanistic model prediction of the baculovirus passage distribution in the infection reactor at steady-state as a function of the residence time. The reported results represent the fraction of passage *N* (i.e., the passage supplied in the feed) among (i) total virions in the supernatant and (ii) viral genomes in the nuclei of infected cells. The two predictions are numerically very similar, and their curves overlap. A fixed feed of 2.5 million uninfected cells/mL and 2 PFU/cell is considered in the model simulation.

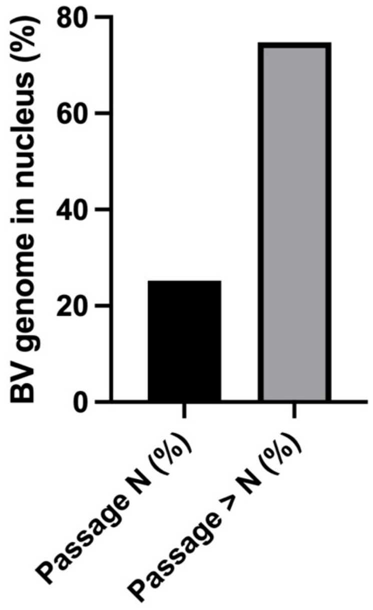


**Figure S3.** Mechanistic model prediction of the fraction of passage $N$ (i.e., the passage supplied in the feed) among (i) total virions in the supernatant and (ii) viral genomes in the nuclei of infected cells in the infection reactor at steady-state. The two predictions are numerically very similar, and their curves overlap. Simulation conditions: 60-hour residence time and feed of 2.5 million uninfected cells/mL and 2 PFU/cell.

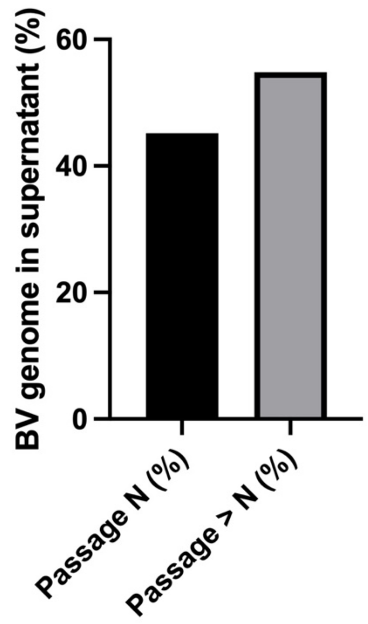

**Figure S4.** Mechanistic model prediction of the fraction of passage $N$ (i.e., the passage supplied in the feed) among (i) total virions in the supernatant and (ii) viral genomes in the nuclei of infected cells in the infection reactor at steady-state. The two predictions are numerically very similar, and their curves overlap. Simulation conditions: 16-hour residence time and feed of 2.5 million uninfected cells/mL and 1 PFU/cell.

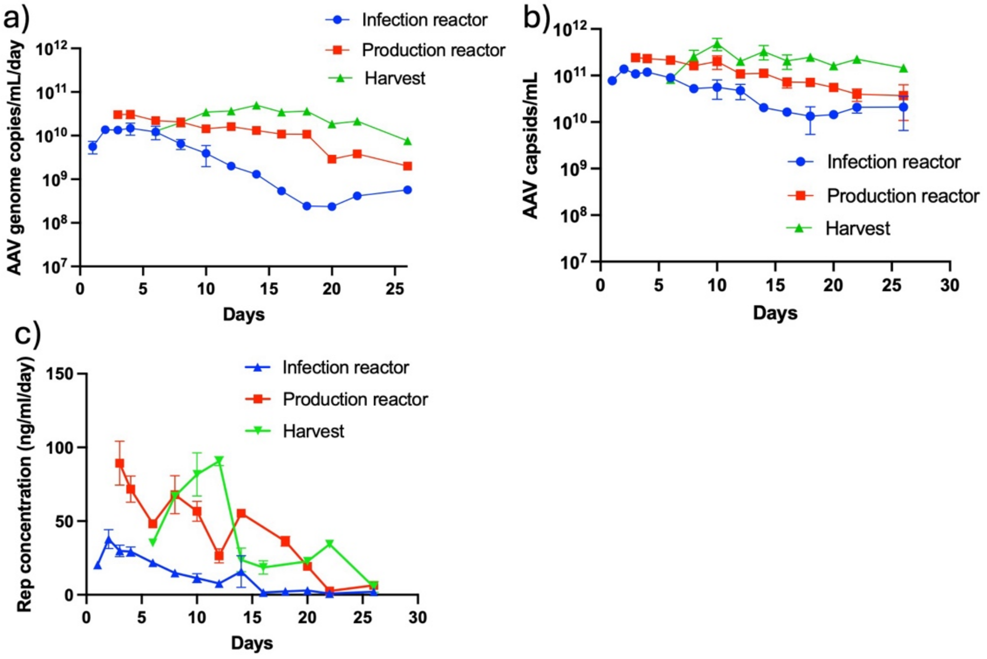


**Figure S5.** Continuous rAAV production in a three-tank cascade without rBV feed. a) and b) total AAV genome and capsid titers were determined from the cell suspension using ddPCR and ELISA, c) Total Rep protein (Rep72 and Rep52) concentrations were obtained from the cell suspension using ELISA. An average residence time of 60 hours was maintained across all three reactors to provide adequate time to self-replicate insect cells and rBV in the growth and infection reactors, respectively. The maximum rAAV yield was achieved in the production reactor.

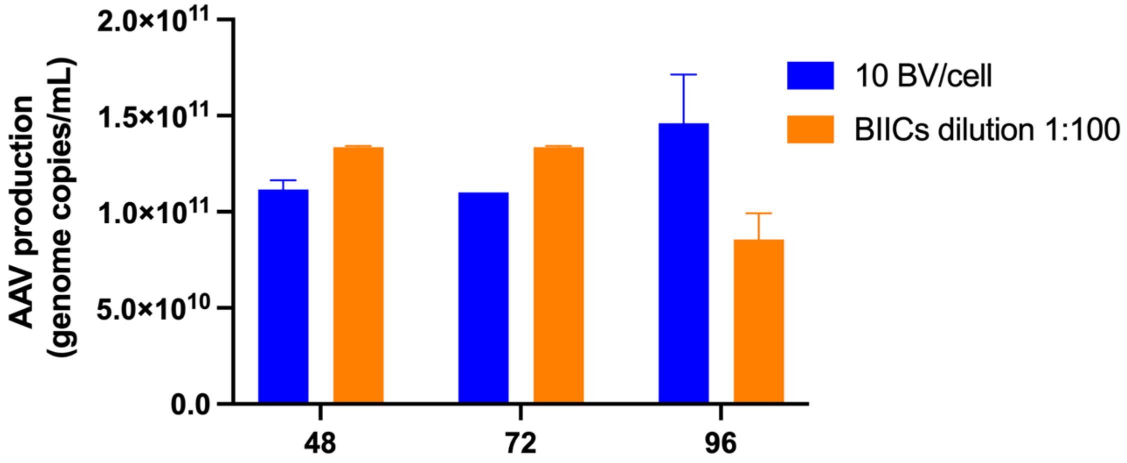


**Figure S6.** Comparison of budded rBV with BIICs for rAAV production. Cell lysates were collected at 48, 72, and 96 hours post-infection. rAAV genome titers were quantified after DNase treatment using ddPCR.
